# Genomic approach to studying the relationship between the CRISPR/Cas system and multidrug resistance in clinical isolates of *Klebsiella pneumoniae*

**DOI:** 10.1101/2024.09.11.612500

**Authors:** Hekmat A. Owaid, Mushtak T.S. Al-Ouqaili, Farah Al-Marzooq

**Author notes:** Corresponding author: Prof. Mushtak T.S. Al-Ouqaili, Department of Microbiology, College of Medicine, University of Anbar, Anbar Governorate, Ramadi, Iraq.

## Abstract

**Background:** Gene editing techniques have been identified as potential tools to combat antimicrobial resistance (AMR). Clustered Regularly Interspaced Short Palindromic Repeats (CRISPR) and associated sequences (Cas) can be key players in this process. Their presence in bacteria can impact the success of this technology in combating AMR. The aim of this study is to investigate whether CRISPR loci are associated by multidrug, extensive drug, or pan-drug resistance in *Klebsiella pneumoniae* from Iraq.

**Methods:** Antibiotic susceptibility testing was performed for 100 isolates to detect patterns of resistance. PCR was used to investigate CRISPR/Cas systems. Whole genome sequencing (WGS) was performed on relevant isolates using a DNA nanoball sequencing platform.

**Results:** Out of 81 *K. pneumoniae* isolates, 81% were resistant to antibiotics, with 71% producing ESBLs and 21% producing carbapenemases. Additionally, 53% were MDR, 19% XDR, and 9% PDR. Complete CRISPR/Cas systems were found in 38% of isolates, while 78% had incomplete systems. Furthermore, intact CRISPR-1, CRISPR2, and CRISPR3 types were found in 27.0%, 34%, and 18.0% of the isolates, respectively. An inverse correlation was found between antibiotic resistance levels and the presence of CRISPR/Cas systems. Two carbapenemase-producing *K. pneumoniae* (XDR and PDR) isolates were characterized by WGS. They were found to be carrying blaNDM-5 and blaOXA-181 genes with additional resistance genes against various antibiotic classes, in addition to CRISPR/Cas systems. Phylogenetic analysis indicated relationships with United Kingdom and Chinese strains. Furthermore, the entire genome revealed the presence of unique virulence and antibiotic resistance genes in *Klebsiella pneumoniae*.

**Conclusion:** An inverse relationship was found between CRISPR/Cas systems and antimicrobial resistance in *K. pneumoniae.* The discovery of blaNDM-5 and blaOXA-181 genes in Iraqi strains is alarming as this can increase the risk of nosocomial outbreaks. This study elucidates the importance of monitoring CRISPR/Cas systems and antimicrobial resistance for more efficient control and prevention of infection in healthcare settings.

## Introduction

The rise of antimicrobial resistance (AMR) across the world is extremely alarming, as highlighted by the very high incidence of superbugs, which are bacteria that have become resistant to various forms of antibiotic (Aslam et al., 2021). *Klebsiella pneumoniae* is one of the illnesses caused by superbugs, and are gram-negative bacteria belonging to the Enterobacterales. This bacterium is particularly infamous for being highly resistant to various antibiotics, with some of its strains considered extensively drug-resistant (XDR) or pan-drug-resistant (PDR) and that can lead to severe life-threatening infections (Sleiman et al., 2021). Chinemerem Nwobodo and colleagues found that *K. pneumoniae* is capable of harboring several genes associated with AMR which are also presented in all other members of the family Enterobacteriaceae, causing high levels of resistance to several forms of antibiotic (Chinemerem Nwobodo et al., 2022). However, one should note that AMR involves carriage of self-transmissible plasmids that facilitate the spread of resistance among bacteria.

AMR in *K. pneumoniae* is mainly due to the creation of powerful enzymes that have the ability to break commonly used antimicrobials belonging to the β-lactam group. These enzymes include carbapenemases and extended spectrum β-lactamase (ESBLs) enzyme production (Oliveira et al., 2022). There are many variants of these enzymes that show high hydrolytic activity against carbapenems, monobactams, and cephalosporins that are commonly used to treat infection (De Angelis et al., 2020). A recent study by Al-Ouqaili et al. (2020) revealed that an increase in the prevalence of carbapenemase-producing *K. pneumoniae*, thereby requiring the use of tigecycline and colistin, which were identified as the only suitable forms of treatment. Among the carbapenemases, IMP, VIM, NDM, KPC, and OXA are the most commonly identified worldwide (Pitout et al., 2020). The latter enzyme has four types, one of which is OXA-48, which is a member of group III. Production of these enzymes by bacteria render them resistant to inhibition by tazobactam, clavulanic acid, and sulbactam (Findlay et al., 2017). Though these enzymes have low affinities to carbapenems, they show a high degree of hydrolytic action towards the penicillins, and their preference for imipenem is highly apparent. A previous study by Mairi et al. (2018) showed that *Enterobacterales* isolates producing OXA-48-like carbapenemase could be identified in human, animal, food item, and environmental samples. To date, there are forty-eight known subtypes of OXA-48-like enzymes, with OXA-181 ranked second subtype (Naha et al., 2021).

Clustered Regularly Interspaced Short Palindromic Repeats (CRISPR) and associated sequences (Cas) protect against invasion of foreign nucleic acid, such as resistance genes and plasmids in several bacterial species (Wu et al., 2021). A study by Wang et al. (2020) showed that the CRISPR-Cas system contains Cas genes sited upstream, spacer sequences derived from different sources, and palindromic repeats. The spacers are the result of incorporation of exogenous DNA from plasmids and phages; after being incorporated into bacterial genome, these spacers help to protect the bacteria from foreign attack with the help of Cas proteins, which can digest sequences in the invader that are identical to sequences in the spacer within bacterial CRISPR arrays (Gholizadeh et al., 2020). It is noteworthy that a recent study reported an inverse relation between the presence of such systems in bacteria and the carriage of AMR genes in multiple bacterial species including *K. pneumoniae* (Wang et al., 2020), but these results are somewhat in conflict with other reports (Tao et al., 2022). As such, there is a demand for more research in this area.

Deep knowledge of the mechanisms of AMR in bacteria is required to design effective strategies to fight resistance. Hence, this study was designed to explore the correlation between these resistance mechanisms in resistant *K. pneumoniae*, shown to be MDR, XDR, and PDR. We also aimed to use whole genome sequencing to find gene variants associated with these systems in selected strains with high rates of resistance.

## Patients and methods

### Ethics statement

The research carried out at the University of Anbar received clearance from the Committee of Medical Ethics with the designation of approval number 14 on March 15, 2022, in accordance with the Declaration of Helsinki. Written informed consent was obtained from all study participants, which included both patients and their parents.

### The patients and design of the study

This study is a cross-sectional survey that was conducted between April and December 2022 at the Ramadi Teaching Hospitals. By employing bacteriological diagnostics on a wide variety of patient clinical specimens, the researchers were able to identify 100 distinct strains of *K. pneumoniae*. A small percentage of the ear swabs (6%), a larger percentage (17%, 50%), and another large percentage (27%), were taken from patients who had had catheterization, otitis media, osteomyelitis, or diabetic foot infections, respectively.

### Bacteriological identification

The hospitals’ microbiology laboratories carried out the handling and cultivation of clinical specimens, along with performing bacteriological studies and confirming biochemical testing. The researchers utilized specialized culture media such as Eosin Methylene blue agar, Blood agar, and MacConkey agar from Germany’s Merck Co. to cultivate all specimens. Subsequently, the specimens were incubated for 24 to 48 hours at 37°C. In addition, the study identified *K. pneumoniae* isolates by examining their cultural and morphological characteristics, such as their growth on lysine iron agar, triple sugar iron agar, their ability to utilize urea and citrate, their production of ornithine decarboxylase, and their appearance under a gram stain. The bacterium was definitively identified utilizing VITEK®2 Compact B System’s GN ID cards, both manufactured by BioMérieux, France (Saki et al., 2022; Oliveira et al., 2022).

### Examination for microbial resistance

To determine the isolates’ sensitivity to antibiotics, the VITEK®2 Compact B System’s AST-GN cards were utilized according to the manufacturer’s guidelines. The results for susceptibility testing for antimicrobials (AST) were obtained for the following antibiotics: levofloxacin, ciprofloxacin, amikacin, gentamicin, cefoxitin, ceftazidime, cefepime, ceftriaxone, cefazolin, ertapenem, imipenem, tigecycline, and nitrofurantoin.

According to the criteria set by the CLSI, results are reported using MIC values. According to the recommendation made by Kurumisawa et al. (2021), the samples were categorized as either resistant, intermediate, or susceptible. According to Gaur et al. (2023), the EUCAST set criteria were used to report tigecycline AST results in 2021. A cross-check of the antimicrobial susceptibility test was performed via an internal quality control using *E. coli* ATCC 25922.

Analyzing the bacteria’s antibiotic susceptibility profiles indicated that these bacteria were multidrug-resistant (MDR), meaning that they were resistant to at least three different groups of antimicrobial agents. Any strain having resistance against not less than six antibiotics was considered to be XDR. The last criterion by which one can consider an isolate to be PDR is its resistance to all the antibiotics examined (Al-Hasani, 2023).

## Molecular assay

### DNA extraction

Genomic DNA was extracted according to the guidelines provided by the manufacturer using the SaMag-12TMDNA extraction machine and SaMag bacterial DNA extraction kit (SaMag, Italy). The suitability of the DNA samples for future use was evaluated via a Quantus^TM^ fluorometer (Promega, USA) (Hussein et al., 2021).

### Detection of CRISPR/Cas genes

As indicated by Wang et al. (2020), the CRISPR/Cas genes were investigated and amplified using specific primers by the conventional polymerase chain reaction (PCR), as reported in Table 1. The Cas1 and Cas3 genes were treated with an initial denaturation temperature of 95°C for 5 minutes. The next step in the denaturation was performed at 94°C for 1 minute to unfold the target gene; this was followed by annealing at a temperature of 60°C for 30 seconds. Finally, extension was performed at a temperature of 72°C for 1 minute for every cycle of the polymerase chain reaction (PCR). The denaturation, annealing, and extension procedure, which consists of 35 cycles, was carried out. The final stage of extension performed for 10 minutes at a temperature of 72°C. The PCR cycle for all CRISPR types involves the following steps: the PCR starts with initial denaturation of the DNA at 95°C for 5 minutes, followed by 1 minute of denaturation at 94°C, 1 minute of annealing at 63°C, and 1 minute of extension at 72°C. This was then denatured, annealed, and extended over 35 cycles, with the final extension phase lasting for 10 minutes at a temperature of 72°C.

Gel electrophoresis was then performed using a 1.5% agarose gel that included 5% ethidium bromide for separating the PCR products. The electrophoresis was carried out using 1X TBE buffer at 50 V for 5 minutes, followed by 100 V for 1 hour. To confirm the appropriate size of the PCR product band, we conducted a comparison with a DNA ladder containing 100-base pair fragments. The UV transilluminator was utilized to identify the fluorescent band following the completion of agarose gel electrophoresis.

### The detection of the CRISPR/Cas system

To determine the presence of CRISPR and its related Cas proteins, the following primers were utilized in PCR:

**Table 1:**
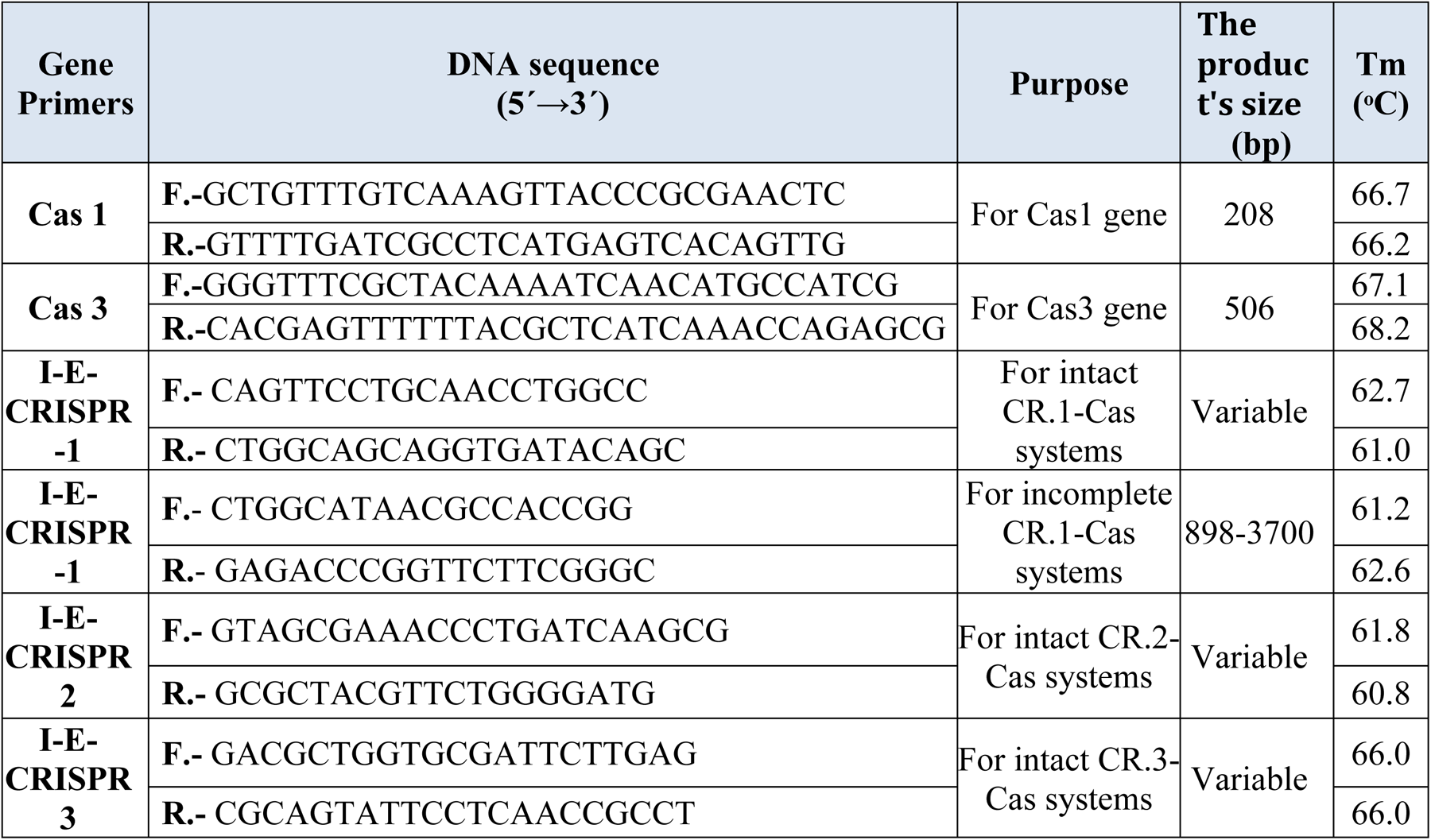
Detailed information about gene primers, including their names, sequences, sizes, guanine:cytosine ratio, melting point, and intended use.

### Whole genome sequencing (WGS)

WGS was used for the two selected clinical isolates of *K. pneumoniae.* The DNA nanoball sequencing platform from BGI-Tech (Hong Kong) was used. Concentrations were measured using a Qubit Fluorometer (Invitrogen). The purities and integrities of the samples were assessed using agarose gel electrophoresis, with a concentration of 1% agarose gel, a voltage of 150 V, and an electrophoresis time of 40 min. Covaris randomly fragmented 1 μg of genomic DNA. The genomic DNA fragments were carefully selected so as to have an average size of 200-400 bp using an Agencourt AMPure XP-Medium kit. After being end-repaired, the fragments were 3’ adenylated, and the 3’ adenylated segments had adaptors ligated to their ends. The adaptor-containing fragments from the prior stage were amplified in this manner. The Agencourt AMPure XP-Medium kit was used to purify the PCR products. After being heat-denatured, the double-stranded PCR products were circularized via the splint oligo sequence. Finally, ssCir DNA, or single-strand circular DNA, was employed as the library format. The library passed the quality control test. The libraries that met the criteria were sequenced using the BGISEQ-500 system. The DNA nanoballs (DNBs) that were created via rolling-cycle replication included more than 300 copies of the reference material. A DNA nanochip with high density was used to load the DNBs into the patterned nanoarray. The final step was to use combinatorial anchor synthesis (cPAS) to get 100 bp reads on both ends.

### Bioinformatics’ analysis

Genome assembly and annotation were achieved using the bacterial bioinformatics resource center (https://www.bv-brc.org/), which was also applied to the PATRIC phylogenetic study. The report on Comprehensive Genome Analysis includes a phylogenetic analysis using the reference and representative genomes provided by PATRIC. To determine the phylogenetic position of the genome, the PATRIC global protein families (PGFams) were selected from the closest reference and representative genomes identified using Mash/MinHash. Following the alignment of protein sequences utilizing muscle, the nucleotides that corresponded to each sequence in these families were accurately assigned to the protein alignment. RaxML was used to analyze the merged sets of amino acid and nucleotide alignments in a single data matrix. The phylogenetic tree’s support values were generated using fast bootstrapping.

The contigs were uploaded to Pathogen Watch (https://pathogen.watch) to confirm species identification and to detect antibiotic resistance genes. Virulence genes were analyzed using Abricate (https://github.com/tseemann/abricate). The PATRIC Genome Annotation Service employs a k-mer-based approach to detect AMR genes. This technique utilizes the well-selected set of typical AMR gene sequence variants that PATRIC maintains. It assigns a functional annotation, the drug class, the broad antibiotic resistance mechanism and, in some circumstances, the specific antibiotic to which the AMR gene confers resistance. The genome maps of the KP-1 and KP-2 strains were visualized by Proksee (Daoud et al., 2023).

### Statistical analysis

The data was examined utilizing the percentage, mean, and frequency methodologies. SPSS 20.0, a statistical analysis program, was utilized to assess the statistical significance of the associations, with a threshold P-value of 0.05. The study’s statistical analysis involved the use of chi-squared analysis and Fisher’s exact test.

## Results

### Susceptibility to antimicrobial agents

One hundred different *K. pneumoniae* strains were isolated from different clinical samples for the investigation. All of the isolates were tested for their susceptibility to various antibiotics, the findings of which are shown in Figure 1. The study found that the bacteria were highly resistant to a variety of antibiotics that target specific bacterial processes, including cell wall construction, DNA synthesis, protein synthesis, and permeability.

Treatment with β-lactam antibiotics, including β-lactam/β-lactamase and carbapenems inhibitors, was successful for the majority of isolates. Aminoglycosides and fluoroquinolones are two examples of other drugs that have shown resistance. Figure 1 shows that the majority of bacteria exhibited resistance to cefazolin, ceftazidime, ceftriaxone, and cefepime, while only a

**Figure 1:**
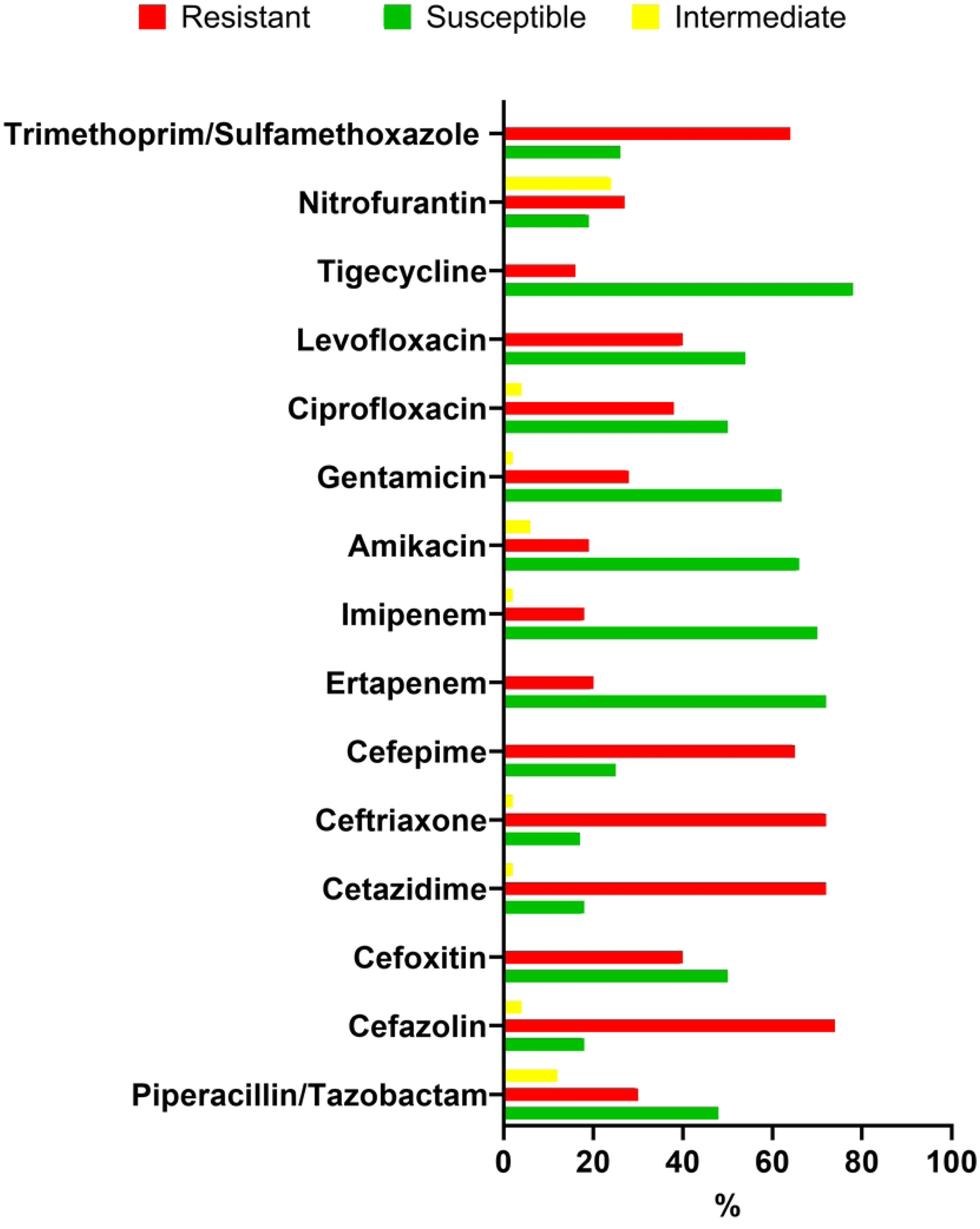
The percentages (MICs) representing the antibiotic resistance patterns of *K. pneumoniae* strains. (In the figure adjust sensitive to Susceptible.

### Identification of CRISPR loci:)

The Cas1, Cas3, CRISPR-1, -2, and -3 genes were identified via PCR. Our analysis revealed comparable patterns in the 100 isolates, which were dependent on the presence of Cas and/or CRISPR genes. Initially, we analyzed the selected *K. pneumoniae* isolates to determine if they had type I-E CRISPR-1 genes. Due to the critical significance of the Cas1 gene in the CRISPR‒Cas system, we utilized primer sequences that were particularly designed to target this specific gene. Based on Figure 2, certain isolates exclusively possessed either CRISPR or Cas, but others contained both system components. The system was detected in 38% of the isolates examined, and we observed a range of gene combinations. Significantly, 53% of the isolates exhibited either CRISPR or Cas exclusively, without any other components. It is worth noting that only nine isolates, accounting for 9% of the total, did not possess this system and the corresponding Cas protein, as depicted in Figure 2.

**Figure 2.**
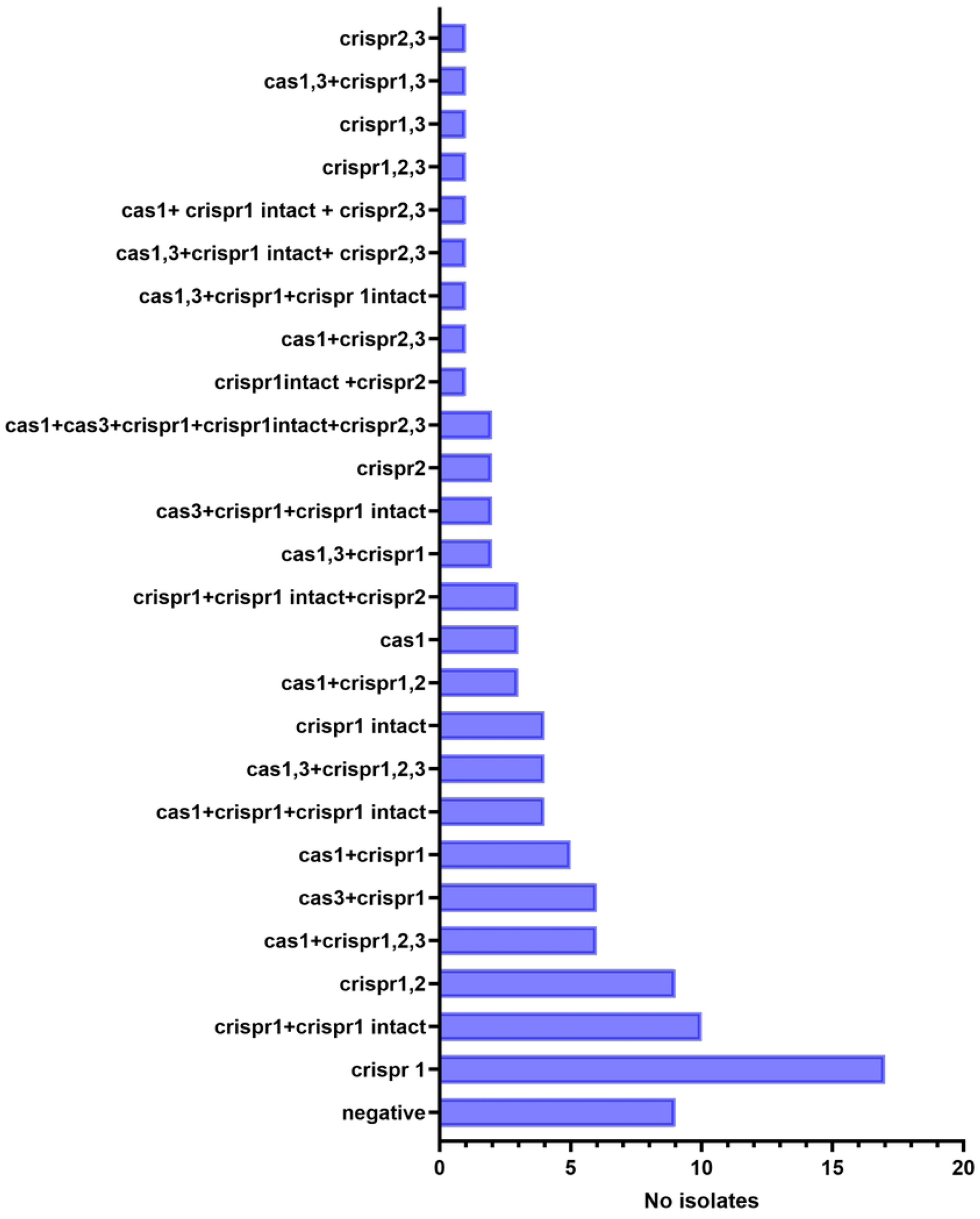
The identification of CRISPR and the genes that encode Cas that are linked to it is particularly connected to the PCR results. The count of isolates possessing either one or both of the genes is displayed, together with the count of isolates without these genes.

*K. pneumoniae* isolates with Cas1 and Cas3 components were identified in 32% and 18% of the samples, respectively. About 78% of the isolates had the primer I-E CRISPR-1 incomplete, which is found on both ends of the *K. pneumoniae* CRISPR-1 gene (Essoh et al., 2013). Also, 27% of the *K. pneumoniae* isolates harbored IE-CRISPR-1, which, in full CRISPR-Cas systems, exhibits several bands corresponding to distinct gene spacers. In a similar vein, IE-CRISPR2 was found in 34% of the *K. pneumoniae* isolates and IE-CRISPR3 in 18%; the bands observed in these genes varied according to the spacers within them, as shown in Figure 3.

**Figure 3:**
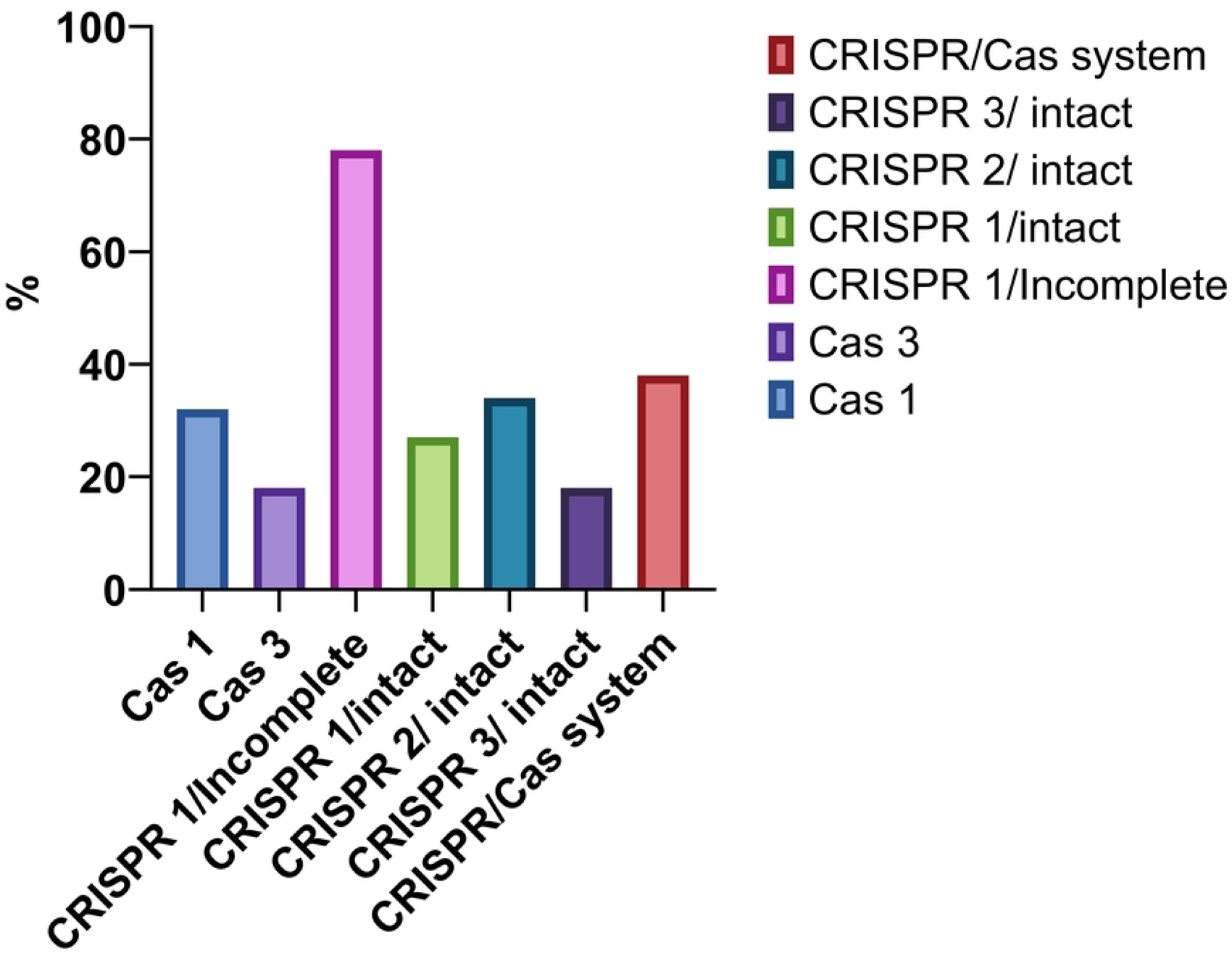
Distribution of the CRISPR Cas elements among the *K. pneumoniae* isolates.

### The CRISPR and Cas protein-associated nosocomial *K. pneumoniae* strains and their associations with antibiotic resistance

Among the isolates tested, 71% were resistant to several antibiotics and developed ESBL, according to the report. Also, all of the 21% carbapenem-resistant isolates were metallo-β-lactamase (MBL) producers. 32% of the Cas1-positive PCR isolates had a CRISPR array, according to the PCR study. Among the bacteria tested, 38% produced ESBLs and 28.6% were resistant to carbapenems; the majority of these bacteria possessed the CRISPR/Cas system. Based on Figure 4, a negative correlation was observed between the presence of antimicrobial resistance genes and the existence of the CRISPR/Cas system. This suggests that the system may play a role in preventing drug-resistant *K. pneumoniae* from acquiring these genes.

**Figure 4:**
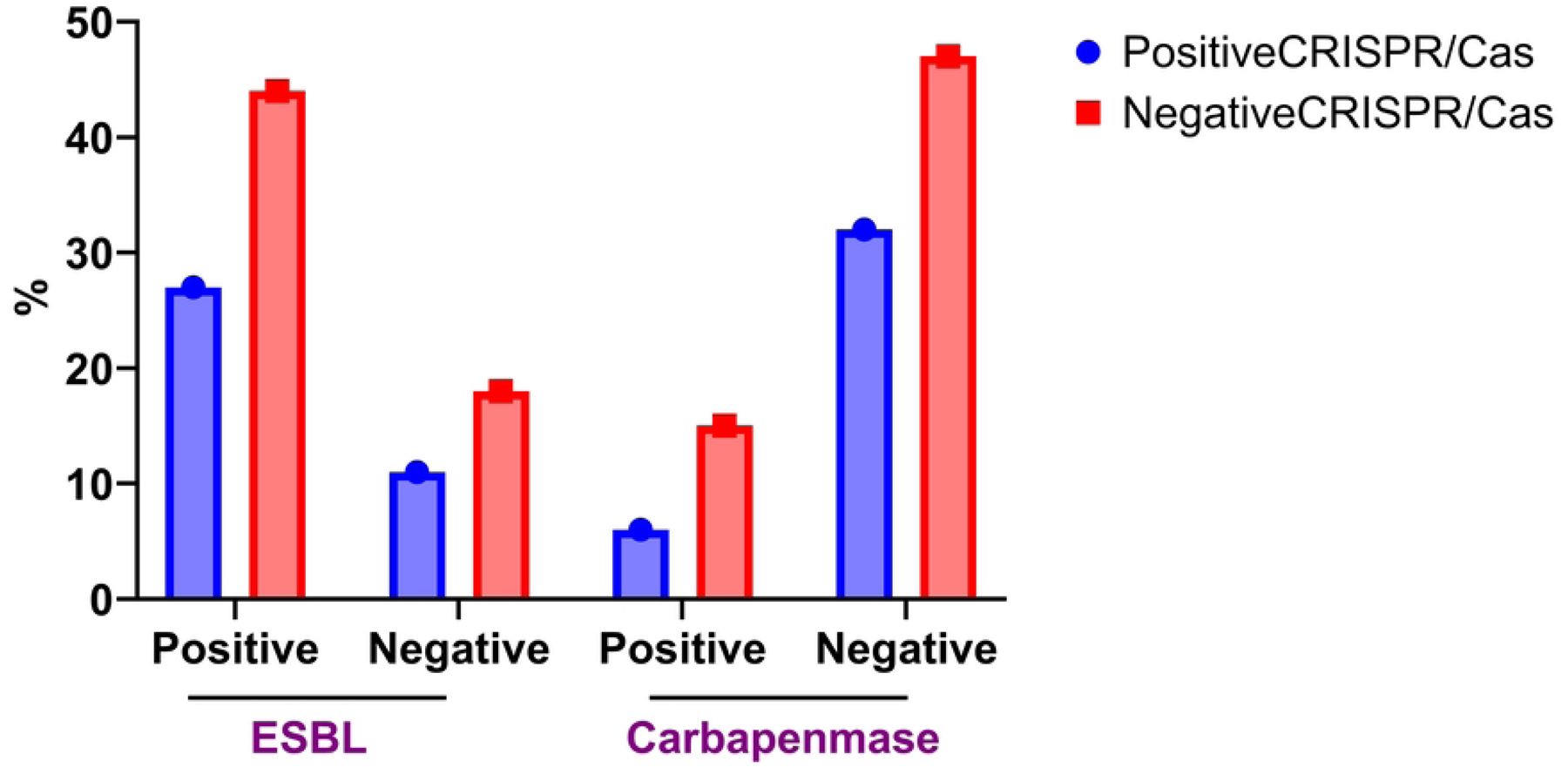
The frequency of the CRISPR/Cas system in *K. pneumoniae* isolates that produce ESBLs and carbapenemases.

### The correlation between the presence of the CRISPR/Cas system and antibiotic resistance

It was found that *K. pneumoniae* isolates absent in the CRISPR/Cas system showed a greater occurrence of antibiotic resistance than those with the system. The subsequent proportions of CRISPR/Cas-resistant isolates were identified for each antibiotic: The following antibiotics have the indicated percentages of effectiveness: piperacillin/tazobactam (10%), ceftazidime (30%), cefepime (27.8%), ceftriaxone (30%), cefoxitin (14.4%), cefazolin (30%), ertapenem (5.6%), imipenem (4.4%), gentamicin (10%), amikacin (5.6%), levofloxacin (15.6%), ciprofloxacin (14.4%), trimethoprim/sulfamethoxazole (26.7%), tigecycline (4.4%) and nitrofurantoin (8.9%). The percentages of CRISPR/Cas-negative isolates that exhibited resistance to various medications were as follows: 62.2%, 23.3%, 50%, 30%, 50%, 50%, 44.4%, 15.6%, 15.6%, 15.6%, 15.6%, 15.6%, 21.1%, 26.7%, 27.8%, 11.1%, 21.1%, and 46.7%, respectively. The data in Figure 5 indicates that resistant isolates lacking the CRISPR/Cas system are more likely not to have CRISPR. However, this disparity is not statistically significant.

**Figure 5:**
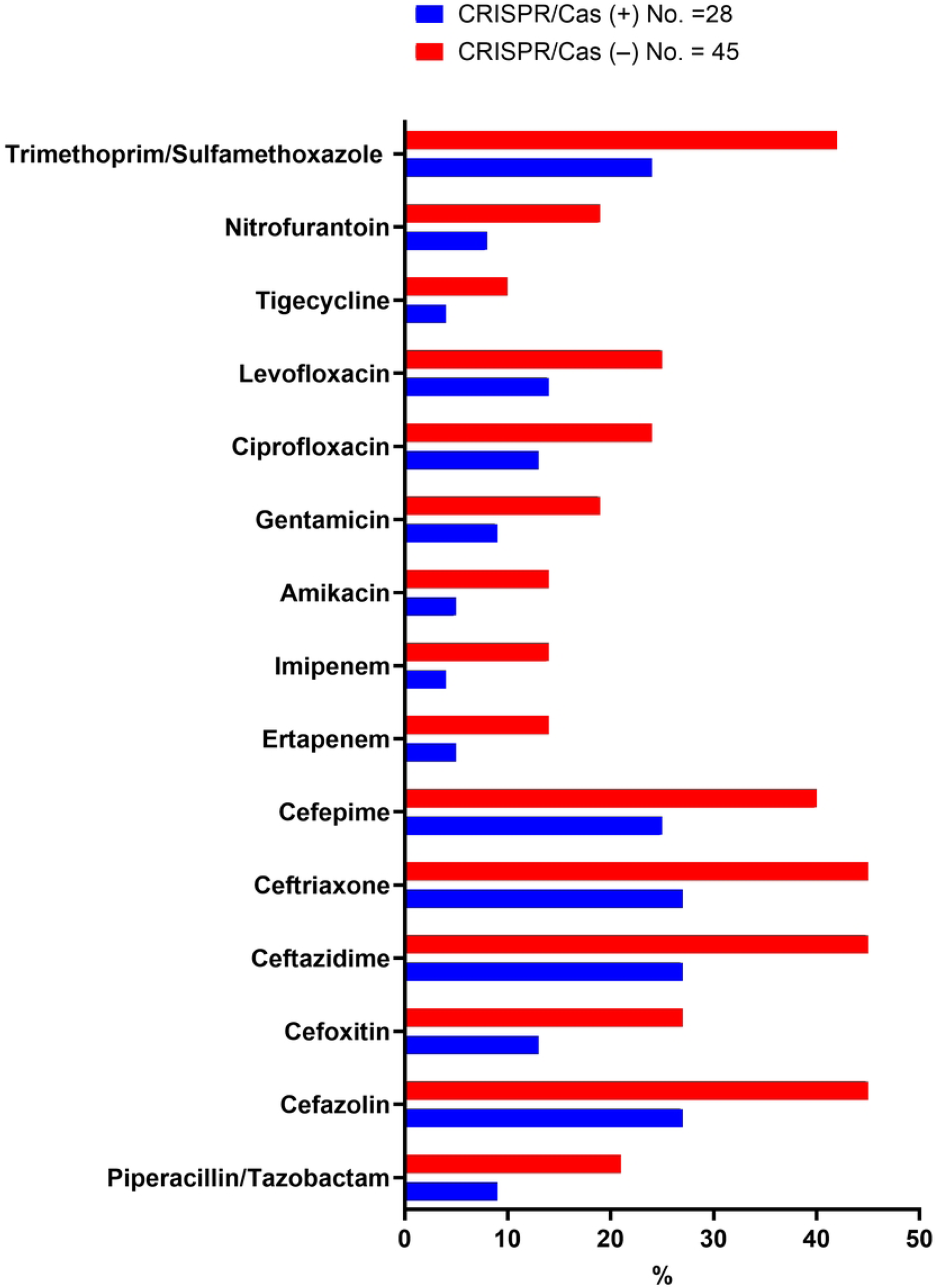
The frequency of positive and negative CRISPR/Cas strains to resistance ratio of chosen antimicrobial drugs. The analysis considered a P-value of less than 0.05 to represent the limit of statistical significance.

Following a functional CRISPR-1 gene, CRISPR-2, and CRISPR-3 were n = 25 (25%, n = 24 (24%), and n = 13 (13%), respectively; our research showed that antibiotic-resistant bacteria are most likely to have a faulty CRISPR-1 gene on both sides. A CRISPR-1 gene that was not entirely functioning (n = 17; 17%), a fully functional CRISPR-1 gene, a CRISPR-2 gene, and a CRISPR3 gene were n = 4 (4%), n = 11 (11%), and n = 5 (5%), respectively, however, were more common in drug-sensitive non-resistant isolates.

The CRISPR/Cas system was found to be inadequate in the vast majority of isolates categorized as XDR or MDR. The majority of MDR isolates (n = 23.0; 43.4%), some resistant XDR isolates (n = 6; 31.5%), and a small percentage of PDR isolates (n = 2.0; 22.2%) contained the CRISPR/Cas system. Nevertheless, upon examining the resistant isolates, which encompassed both XDR and PDR isolates, a substantial quantity of such (n = 20) were discovered to lack the CRISPR system and its associated Cas proteins, as shown in Figure 6.

**Figure 6:**
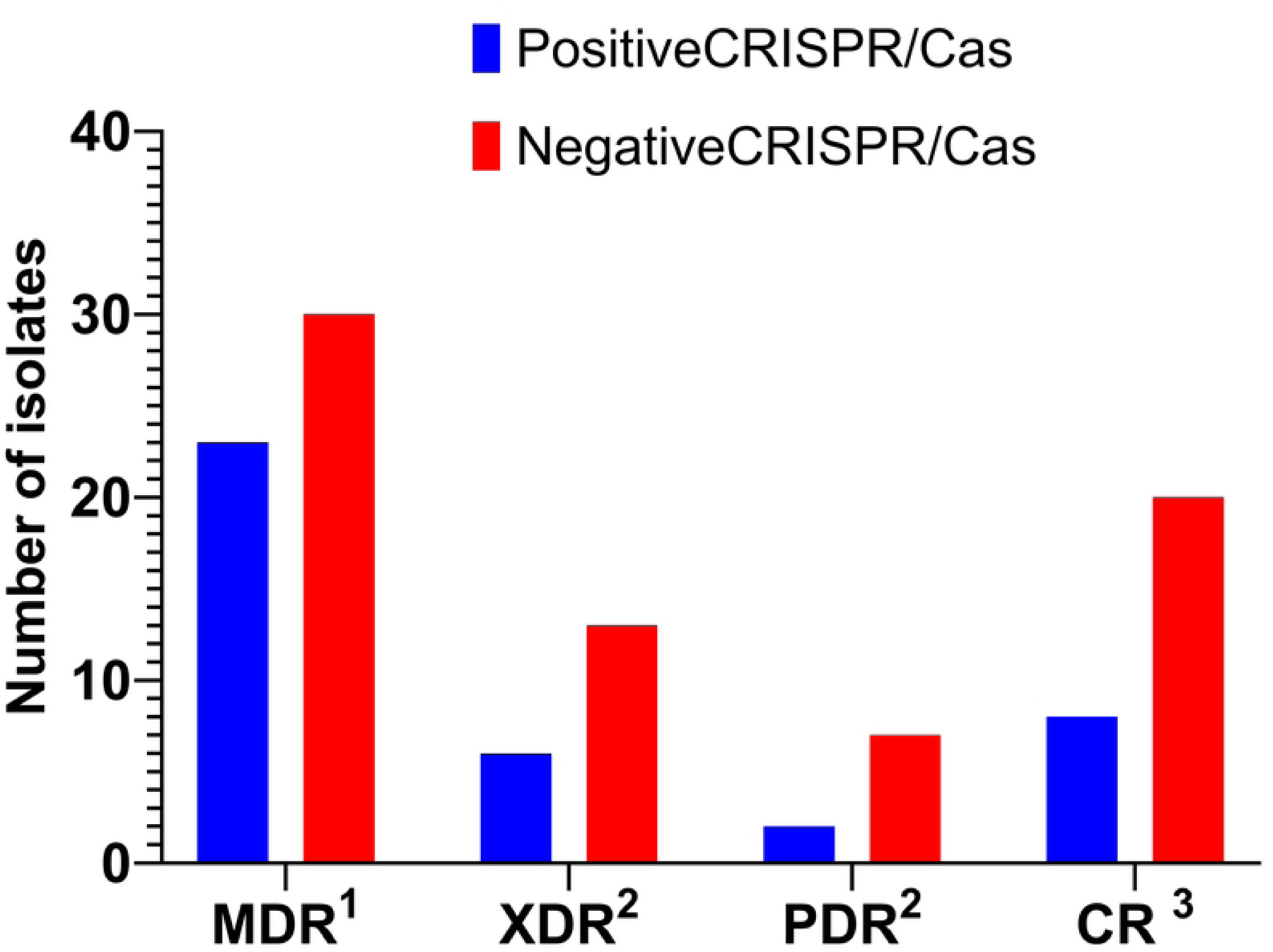
Examining the relationship between the incident rate of the CRISPR/Cas system and MDR, XDR, and PDR. All MDR isolates exhibited the production of ESBLs.^2^ XDR and PDR mostly consisted of metallo-β-lactamase producers.^3^ CRs isolates were a combination of PDR and XDR.

### Fundamental statistics of the genome assembly

The G+C content of *K. pneumonia* (KP-1) is 56.92%, and the genome assembly size is 5,770,550 bp, divided into 97 contigs. The genome contains 5,728 protein-coding sequences, along with an additional 81 sequences that are coding RNAs (including 78 tRNAs and 3 rRNAs). The minimum sequence length in the genome is 295,571 bp, which makes up 50% of the genome, and the L50 count is six, representing the smallest number of contigs whose combined length gives the N50. In contrast, the genome assembly size of *K. pneumonia* (KP-2) is 5,664,670 bp, with a G+C content of 57.13% spread across 106 contigs. The genome contains 84 coding RNAs (81 tRNAs and 3 rRNAs) and 5,666 protein-coding sequences. The genome’s N50 is 218,244 bp, and its L50 count is eight.

The distribution of genome annotations is graphically presented in a circular diagram (Figure 7), including contigs, GC content, GC skew, CDS on the forward and reverse strands, RNA genes, as well as CDS with similarity to known antimicrobial resistance genes. These annotations are displayed in concentric rings, with the exterior rings showcasing the contigs, GC content, and GC skew, progressing towards the interior rings that represent RNA genes, CDS, and related subsystems, which are differentiated by the colors of the CDS on the forward and reverse strands.

**Figure 7.**
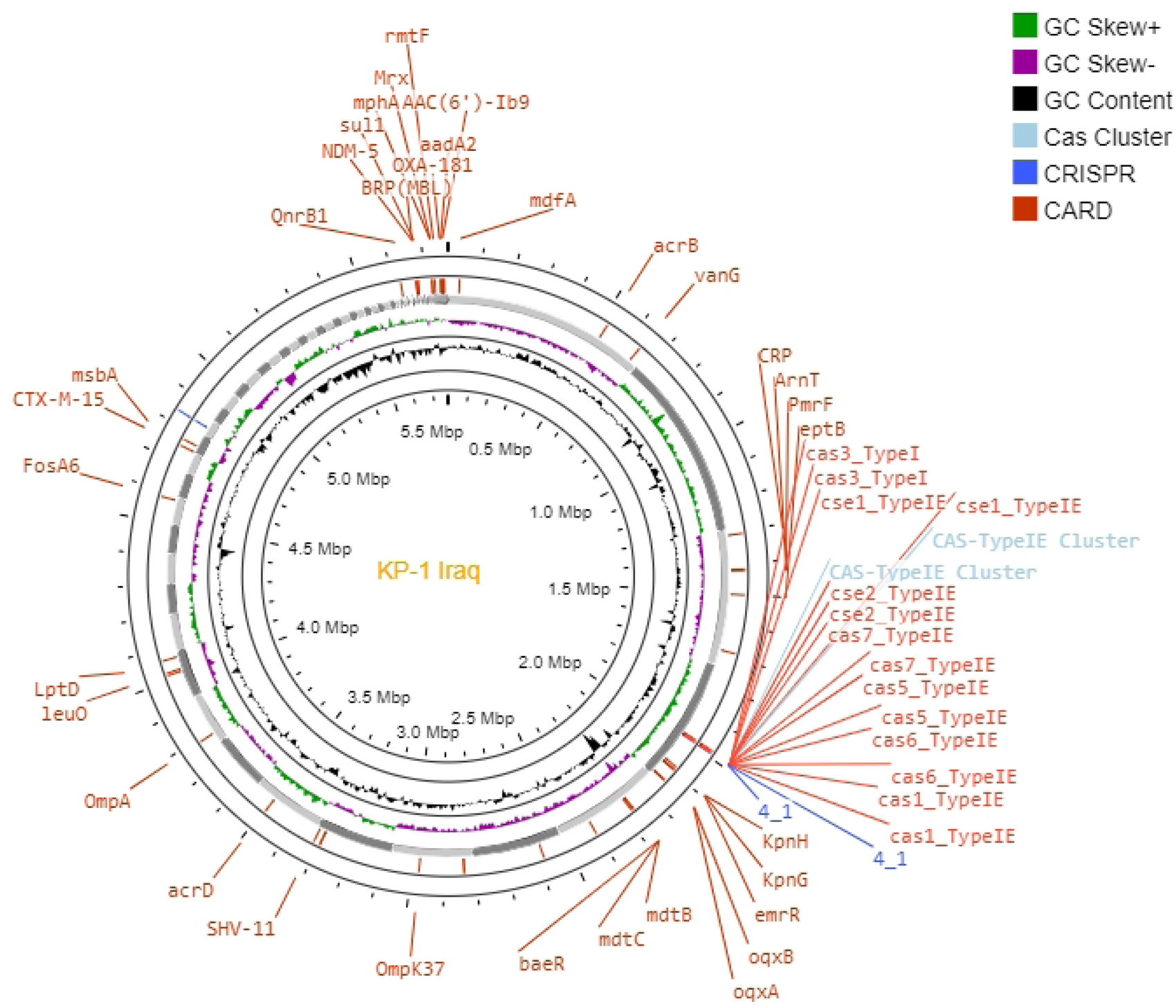
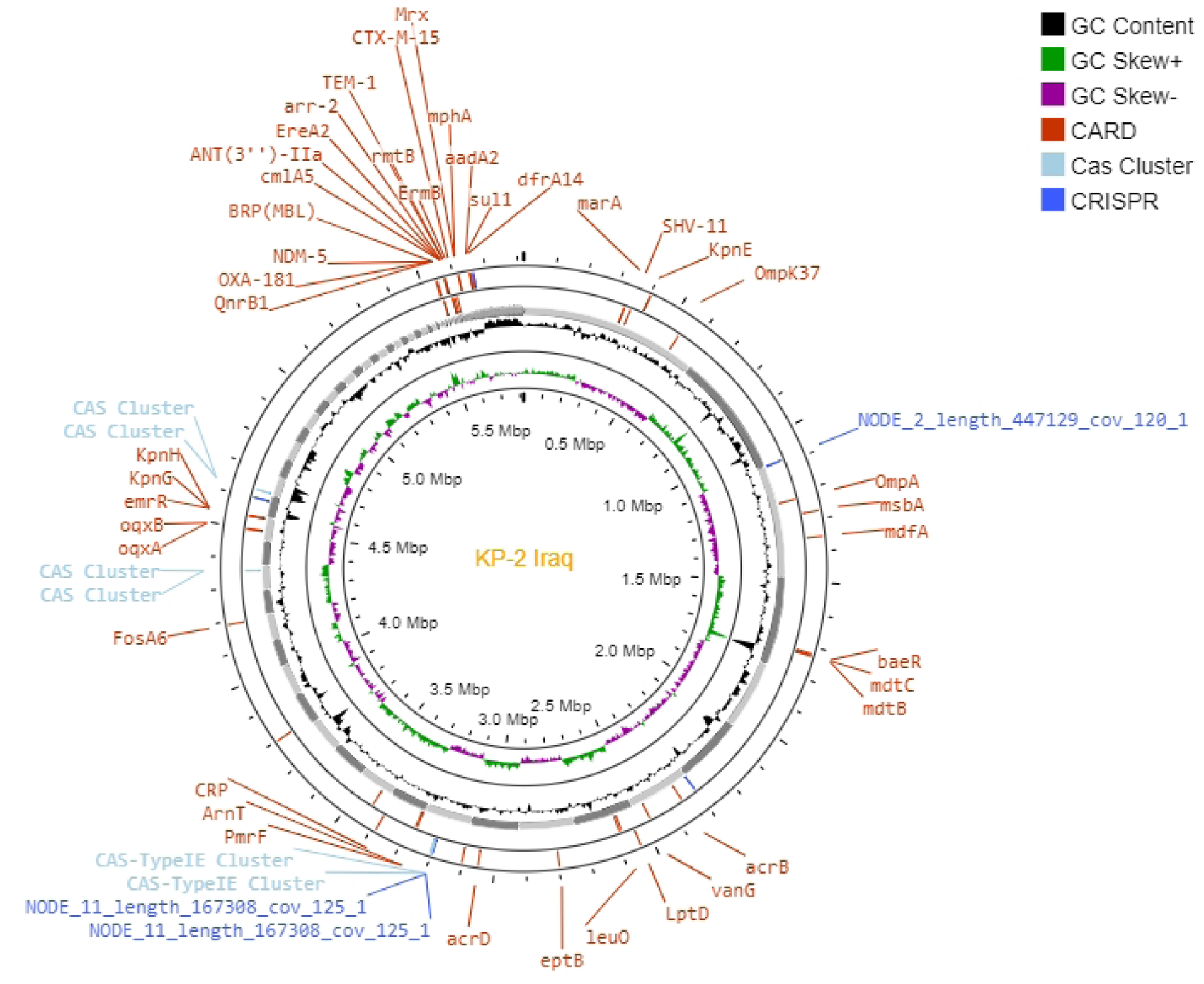
(a and b): Antibiogram, AMR genes and circular genome mapping of clinical isolates of *K. pneumoniae* 1 (KP-1) in part a and *K. pneumoniae* 2 (KP-2) in part b.

### Antimicrobial Resistance (AMR) profiling

Antimicrobial resistance patterns in KP-1 and KP-2 were characterized phenotypically. It was found that both strains have developed resistance to a significant number of antibiotics commonly used in medical care. It is important to consider specific AMR pathways, especially the presence or absence of SNP mutations that indicate resistance. An overview of the AMR genes identified in the genomes, as well as the relevant AMR mechanism, can be found in Table 2.

**Table 2.**
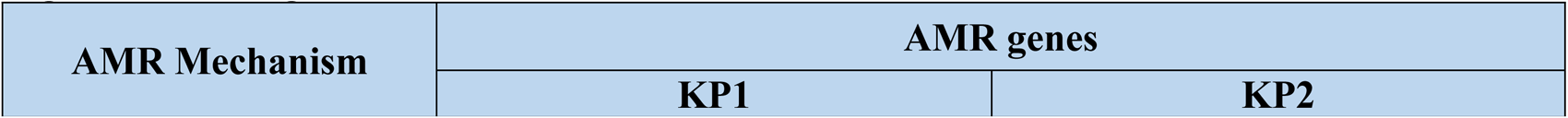
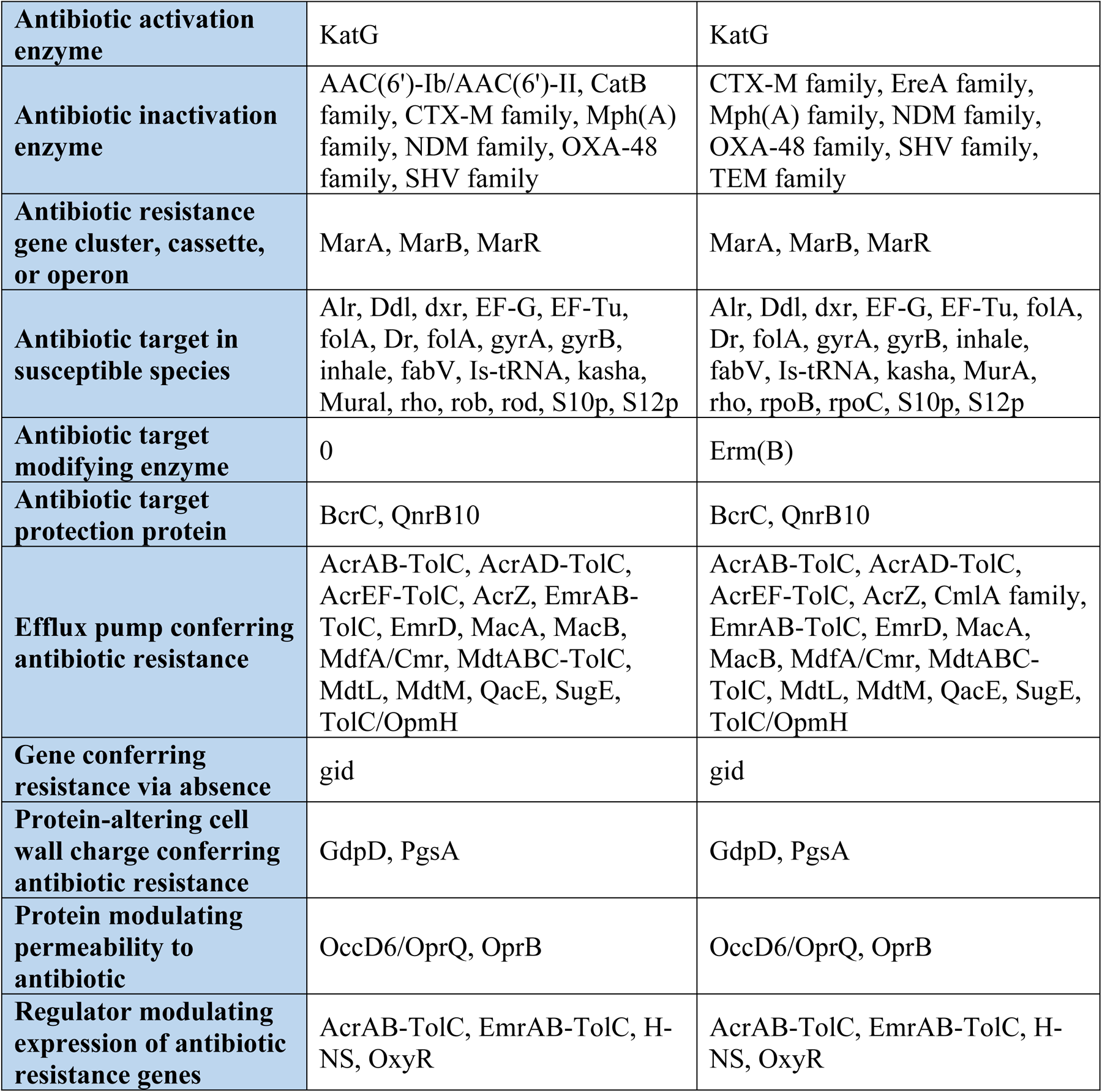
The distribution of sequence variants of antimicrobial-resistant genes in both *K. pneumoniae* 1 (KP-1) and *K. pneumoniae* 2 (KP-2) against the eight groups of antimicrobial agents according to mechanism of action.

| AMR Mechanism | AMR genes |  |
| --- | --- | --- |
|  | KP1 | KP2 |
| **Antibiotic activation enzyme** | KatG | KatG |
| **Antibiotic inactivation enzyme** | AAC(6')-Ib/AAC(6')-II, CatB family, CTX-M family, Mph(A) family, NDM family, OXA-48 family, SHV family | CTX-M family, EreA family, Mph(A) family, NDM family, OXA-48 family, SHV family, TEM family |
| **Antibiotic resistance gene cluster, cassette, or operon** | MarA, MarB, MarR | MarA, MarB, MarR |
| **Antibiotic target in susceptible species** | Alr, Ddl, dxr, EF-G, EF-Tu, folA, Dr, folA, gyrA, gyrB, inhale, fabV, Is-tRNA, kasha, Mural, rho, rob, rod, S10p, S12p | Alr, Ddl, dxr, EF-G, EF-Tu, folA, Dr, folA, gyrA, gyrB, inhale, fabV, Is-tRNA, kasha, MurA, rho, rpoB, rpoC, S10p, S12p |
| **Antibiotic target modifying enzyme** | 0 | Erm(B) |
| **Antibiotic target protection protein** | BcrC, QnrB10 | BcrC, QnrB10 |
| **Efflux pump conferring antibiotic resistance** | AcrAB-TolC, AcrAD-TolC, AcrEF-TolC, AcrZ, EmrAB-TolC, EmrD, MacA, MacB, MdfA/Cmr, MdtABC-TolC, MdtL, MdtM, QacE, SugE, TolC/OpmH | AcrAB-TolC, AcrAD-TolC, AcrEF-TolC, AcrZ, CmlA family, EmrAB-TolC, EmrD, MacA, MacB, MdfA/Cmr, MdtABC-TolC, MdtL, MdtM, QacE, SugE, TolC/OpmH |
| **Gene conferring resistance via absence** | gid | gid |
| **Protein-altering cell wall charge conferring antibiotic resistance** | GdpD, PgsA | GdpD, PgsA |
| **Protein modulating permeability to antibiotic** | OccD6/OprQ, OprB | OccD6/OprQ, OprB |
| **Regulator modulating expression of antibiotic resistance genes** | AcrAB-TolC, EmrAB-TolC, H-NS, OxyR | AcrAB-TolC, EmrAB-TolC, H-NS, OxyR |

### Phylogenetic Analysis

The phylogenetic tree shows the evolutionary relationship between KP-1 and KP-2 in comparison to other *K. pneumonia* strains reported from the same or similar sources across continents in 2021–2023, as depicted in Figure 8.

The phylogenetic analysis revealed that the KP-1 PDR strain is closely related to strains originating from the United Kingdom, while the KP-2 XDR strain shows an evolutionary relationship with strains from China. Additionally, the entire genome analysis indicated the presence of distinctive virulence and antibiotic-resistance genes in *Klebsiella pneumoniae*. Notably, KP-1 and KP-2 are genetically separate and are not clonally related to each other.

**Figure 8:**
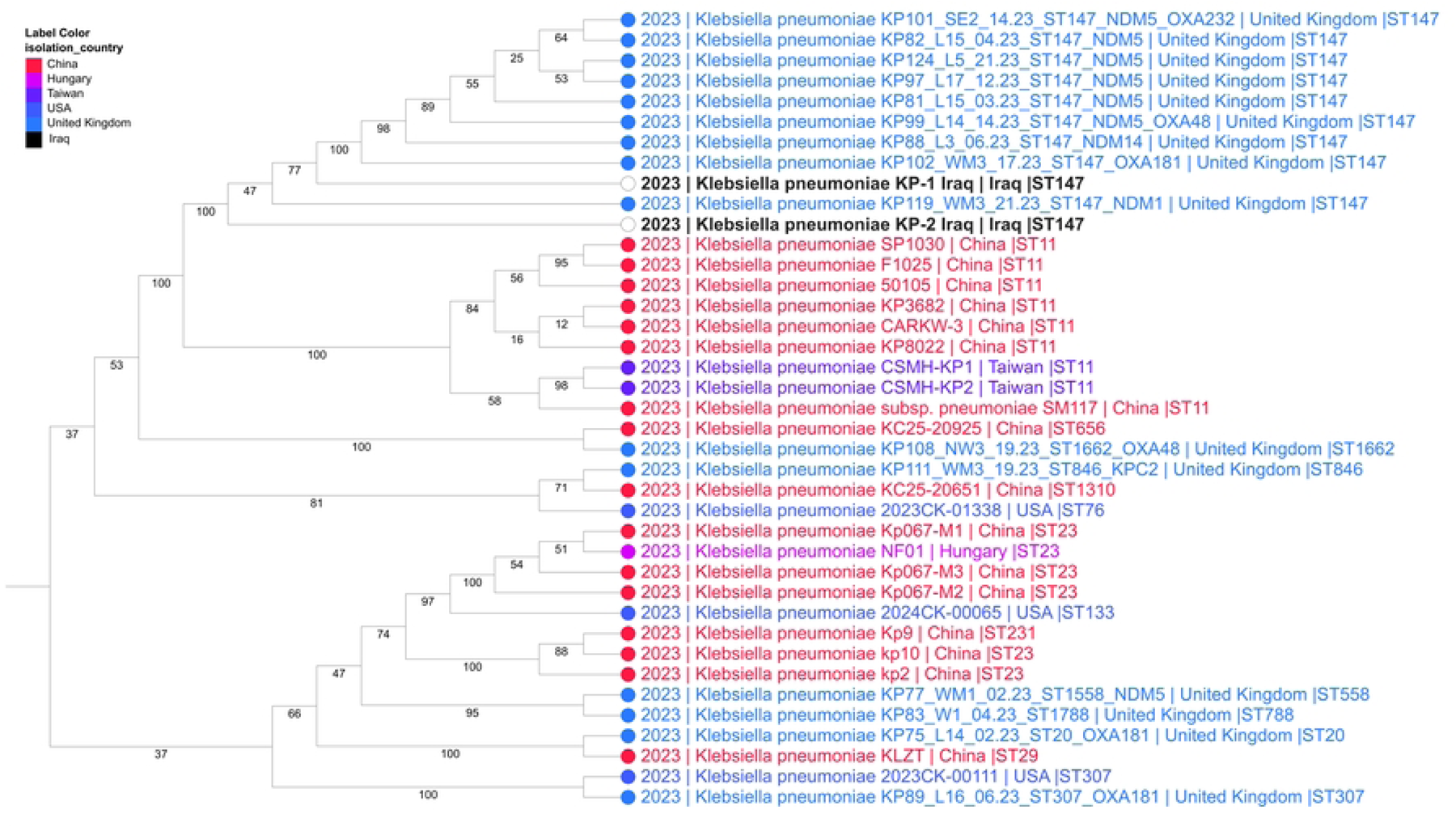
The phylogenetic tree demonstrating the evolutionary relatedness of *K. pneumonia* KP-1 and KP-2 to other strains of the same bacterium that have been subject to global selection. The tree was generated using PATRIC v3.6.2.

## Discussion

Hospitals and other healthcare institutions have been recognized as major sources of the spread of drug-resistant pathogenic bacteria, including the *K. pneumoniae* species (Alduhaidhawi et al., 2022). Bacterial resistance often arises due to the excessive and unregulated use of antimicrobial medicines. The dissemination of antibiotic-resistant microorganisms from healthcare institutions to the wider population represents a significant problem (Samreen et al., 2021).

The majority of penicillins and third-generation cephalosporin medications are ineffective against bacteria that produce ESBLs, primarily *Klebsiella* species and *E. coli*. According to the CDC, approximately 26,000 cases of healthcare-associated infections are attributed to ESBL-producing *Enterobacteriaceae* (Kim et al., 2015). Patients with ESBL-producing isolates have a 57.0% higher risk of death compared to those with non-ESBL isolates (Quirante et al., 2011).

Based on a previous study, *K. pneumoniae* exhibits a remarkably high prevalence of antibiotic resistance to various important antimicrobial drugs (Chaves et al., 2001). The study results revealed a significant level of resistance to β-lactam antibiotics, specifically ceftazidime (73.0%), ceftriaxone (73.0%), and cefazolin (75%). This might be because of misuse and/or abuse of these antimicrobial agents without reference to a medical professional. In their study, Ejaz et al. (2013) found that 88.63% of the *K. pneumoniae* isolates were resistant to third-generation cephalosporin.

Multiple national and international studies have documented a high percentage of such isolates as ranging from moderately to highly resistant. Khalaf and Al-Ouqaili (2018), also showed that the resistance rates to ceftriaxone were 88%, and to ceftazidime, 84%. Al-Kubaisy and colleagues found that 88.2% of *K. pneumoniae* bacteria also showed resistance to ceftriaxone, while 82.3% were resistant to ceftazidime (Al-Ouqaili et al., 2020). In northwest Pakistan, Ullah and colleagues reported a high level of cephalosporin resistance among *K. pneumoniae* isolates (54.35%) that also revealed resistance to ceftriaxone (Ullah et al., 2009). Furthermore, according to Khalaf and Al-Ouqaili (2018), isolates of *K. pneumoniae* were found to be resistant to ceftazidime (92%) and cefotaxime (96%). Resistance to fluoroquinolones has developed in several strains of enterobacteria due to the extensive use of antimicrobial drugs in recent years. Ahmadi et al. (2022) noted that *K. pneumoniae* is more likely to develop resistance to quinolones and other antibiotics. This multidrug-resistant opportunistic bacterium poses a major challenge to medical professionals worldwide with regard to treating infectious diseases. Among the antibiotics tested, ceftriaxone (73.0%) and cefepime (66.0%) were the most frequently used. Among the isolates, 53 (53%) were classified as MDR and 19 (19%) as XDR.

According to the study results, *K. pneumoniae* isolates had a resistance rate of 40.0% to ciprofloxacin and 43.0% to levofloxacin. These findings differed from those reported by Jomehzadeha et al. (2022), who reported resistance rates of 18.5% for ciprofloxacin and 30.4% for levofloxacin. The widespread availability and use of antibiotics, along with issues related to dosage, treatment duration, and inappropriate prescribing, have contributed to a significant increase in antibiotic resistance despite their positive impact on the treatment of infectious diseases (Alhamdany, 2018). The initial proposal was to selectively eliminate certain bacterial genotypes using the CRISPR/Cas system as an antibacterial agent (Bikard et al., 2012).

The objective of this study was to detect the resistance patterns of *K. pneumoniae* to a variety of antibiotics. A total of 100 clinical isolates of *K. pneumoniae* were analyzed, each demonstrating unique resistance patterns. KP-1 and KP-2 were identified as pan-drug and extensively drug-resistant, respectively, based on phenotypic analysis that indicated resistance to multiple drugs, with significant therapeutic implications. Regarding the resistant genes, the results of genomic analysis presented in Table 6 further support these findings. The blaPAO and blaOXA48 family, known as β-lactamase resistance encoding genes, have been documented in multiple studies on the characterization of the *K. pneumoniae* genome (Hussain et al., 2017; Madaha et al., 2020), which is consistent with the strains examined in our study. We observed a unique result in the genome of PDR-KP-1: the detection of the KPC-2 gene. Both strains are dual carbapenemase producers (NDM-5 + OXA-181), which has rarely been reported in our country or, indeed, the Arab nations, but has recently been documented in other parts of the world (Forero-Hurtado et al., 2023). Conversely, the genomic study on XDR-KP-2 revealed the presence of a GES-type ESBL encoding gene which has not been previously reported in Iraq. Furthermore, this strain co-exists with blaPAO, blaOXA-family, and NDM-family carbapenemases, potentially explaining its unique resistance to all the antimicrobial agents tested in our study.

Additional research by Girlich, Naas, and Nordmann (Girlich et al., 2004) suggested that the presence of blaOXA-50 may reduce the susceptibility of *K. pneumoniae* to ticarcillin, ampicillin, moxalactam, and meropenem. Therefore, the presence of blaPAO-, blaOXA-, and NDM-family carbapenemases in the genomes of PDR-*K. Pneumoniae* could explain its resistance to meropenem, colistin, and tigecycline.

According to updated various studies, the CRISPR/Cas system effectively eradicates drug-resistant gene populations from bacterial populations. During this process, plasmids carrying AMR genes can be removed, making bacteria more sensitive to antibiotics. However, before using CRISPR/Cas to target antibiotic resistance in natural microbial communities, several rather significant difficulties must be resolved. Developing a viable distribution plan will be critical to harnessing the full potential of this technology in curbing the spread of AMR due to mobile genomic elements (MGEs), both in ecological and therapeutic contexts. One way to enhance the efficiency of this technique is by reprogramming basic CRISPR/Cas constructs to target certain genes. This approach has the potential to either maintain or restore the antibacterial effectiveness of antibiotics and aid in combating reservoirs of antimicrobial resistance. Demonstrating the precision of the CRISPR/Cas antibacterial mechanism involved isolating specific bacterial strains from a diverse population of *E. coli* genotypes. To achieve this, a plasmid containing the CRISPR/Cas system was introduced. This plasmid was designed to target a specific sequence that is related to each genotype. Additionally, two pieces of research have shown that CRISPR/Cas9 can successfully eradicate *Staphylococcus aureus* and *Escherichia coli* by combining it with phagemids, which are plasmids found within phage capsids. CRISPR/Cas9 constructs were used to target AMR genes through phagemid transduction, successfully eliminating plasmids carrying these genes from bacteria. In a study conducted by Citiotik et al. (2014), CRISPR/Cas9 was altered in bacteria by integrating conjugative plasmids containing chromosomal AMR genes. A further study also confirmed that dangerous bacteria can reduce antibiotic resistance and transit of genes, as well as eliminate the plasmids containing such genes through the introduction of sequence-specific CRISPR/Cas9 genes (Pursey et al., 2018). Our study revealed that our isolates exhibited numerous characteristics of the CRISPR/Cas system. Due to exposure to various bacteriophages throughout their lifespan, CRISPR arrays show significant variation in sequence, size, and quantity. This finding aligns with the observations made by Makarova et al. (2011), who noted that different bacterial species have CRISPR arrays with distinct patterns, lengths, and quantities. The wide variety of cyanophages produced by *Microcystis aeruginosa* supports the findings of Kuno and colleagues, which demonstrated that the DNA of this pathogen contains numerous antiviral defense mechanisms (Kuno et al., 2012).

In a 2014 study by Bondy-Denomy and Davidson, it was found that the synthesis of bacterial virulence factors is linked to CRISPR/Cas systems and defense against the introduction of foreign DNA into these organisms. Although extensive research has been conducted on these systems in various organisms, both harmful and harmless, only a limited number of studies have revealed their actual functioning within a living organism. Locating information on the content of spacers, such as the proportion of spacers that correspond to a known sequence or that are unique to a single species, is challenging in the scientific literature. The composition of the spacers, the ratio of spacers that conform to a recognized sequence, and the spacers unique to a specific species would all be detailed in articles outlining the fundamental workings of this system. The use of this data could help us better understand the actions of these systems in the studied isolates. Additionally, it could help us comprehend how bacteria have evolved and how horizontal gene transfer has impacted both human and environmental health. *K. pneumoniae*, which is among the top five bacteria responsible for hospital-acquired infections worldwide, has not been extensively studied or, indeed, is well understood. It belongs to the ESKAPE group, as noted by Djordjevic et al. (2012). *K. pneumoniae* possesses several ecological benefits such as its large plasmids, virulence factors, and ability to spread easily in hospital wards, which allow it to adapt to different environments (El Fertas-Aissani et al., 2013). Hence, our objective is to ascertain the existence of CRISPR/Cas systems in *K. pneumoniae* and explore their correlation with virulence, resistance to several medicines, horizontal gene transfer, and other characteristics. Scientists have discovered that the CRISPR/Cas system safeguards prokaryotes from viruses by utilizing acquired spacers from infest components to regulate the immune response in a specific sequence (Barrangou and Horvath, 2012). Consequently, the collection of spacers may reflect the bacterial lifestyle (Horvath et al., 2009). Our research indicates that this is the first Arab study to investigate the link between the CRISPR/Cas system and MDR, XDR, and PDR of *K. pneumoniae* strains, as well as WGS. These strains are extremely challenging, if not impossible, to treat using existing antibiotics. Only 38.0% (38/100) of the *K. pneumoniae* isolates tested positive for CRISPR/Cas, suggesting that this percentage is low. Isolates contained CRISPR1/Cas in 38.0% of cases, CRISPR2/Cas in 34.0% of cases, and CRISPR3/Cas in 18.0% of cases. In all likelihood, the low frequency of CRISPR/Cas systems in this study is because the majority of the isolates tested were discovered to be antibiotic resistant. Our findings show that 30.7% of Taiwanese clinical *K. pneumoniae* isolates had the CRISPR/Cas system, in agreement with previous research (Li et al., 2018). According to Wang et al. (2020), only 29 (or 21.32 percent) of the 136 *K. pneumoniae* isolates tested positive for CRISPR/Cas. Only two of the remaining samples were found to be positive for CRISPR3, whereas thirteen tested positive for CRISPR2. We also found that 14 of the isolates contained both genes. Zhou and colleagues found this particular mechanism in approximately one-third of the total 300 K *pneumoniae* isolates they investigated (Zhou et al., 2022). According to the recent study conducted by Lin et al. (2016), out of the total 52 *K. pneumoniae* isolates, only six adopted this particular mechanism. The data reveal that *K. pneumoniae* does not have a highly distributed CRISPR/Cas system among any of its genomes.

This study’s findings are especially compelling since they link the recent development of the CRISPR/Cas system to the absence of antibiotic-resistant genes. According to Ostria-Hernández et al. (2015), the CRISPR/Cas system is not broadly distributed in *Klebsiella pneumoniae*. The study found that the approach worked perfectly with both whole and incomplete genomes. Of the 100 clinical isolates of *K. pneumoniae* acquired from healthcare facilities, 33 (or 33.0%) showed CRISPR/Cas activity. The CRISPR/Cas system was present for between 30.7% (54 out of 176) and 12.4% (27 out of 217) of the population (Lin et al., 2016).

The appearance of antibiotic-resistant bacteria containing CRISPR or Cas genes indicates that these systems were previously widespread but have become non-functional. The elimination of bacteria is necessary to facilitate the acquisition of AMR genes and the subsequent development of resistance (Pinilla-Redondo et al., 2022). Because of the cross-sectional nature of this study, there were few bacteria that were responsive to treatment, with the majority of such causing these disorders being MDR, XDR, or PDR in nature, which is a matter of significant concern. Notably, the PDR bacteria had the lowest frequency of the CRISPR/Cas system, while MDR bacteria had a higher frequency (Jawir et al., 2023); also, those researchers provided evidence that the CRISPR/Cas system is negatively associated with antibiotic resistance. Furthermore, it is important to note that as the level of antimicrobial resistance increases from MDR to XDR to PDR, the frequency of occurrence of the CRISPR/Cas system decreases. This could be because the CRISPR system protects the genome of *K. pneumoniae* from invasion via genetic elements carrying antibiotic-resistant genes, thereby ensuring genetic stability. Moreover, exogenous genetic plasmids and other transferable elements may carry advantageous genes that enhance bacterial fitness, such as virulence and resistance to antimicrobial agents.

Phylogenetic analysis of the relationship between KP-1 and KP-2 with other strains revealed that KP-1 has 100% identity with strains of human origin in the United Kingdom, and shows some phylogenetic proximity to strains from other countries such as China, Taiwan, and the USA. Similarly, the KP-2 strain has 100% identity with strains of human origin in China, and also exhibits some phylogenetic proximity to strains from other countries including the United Kingdom, Taiwan, and the USA. This could be a result of population dynamics, where diseases and genetic material can be transferred between communities when populations relocate. The closer phylogenetic relationship between our isolates and strains from different countries can be explained by increased economic mobility and social contact between the two regions, as demonstrated in Figure 8.

It was concluded that the CRISPR/Cas system and drug resistance have opposing effects. Also, whole genome sequencing of XDR and PDR *K. pneumoniae* revealed that the presence of artefacts such as virulence, CRISPR, and resistance genes contribute to their ability to invade and persist in the host environment, posing a serious threat to public health. The seriousness of the illnesses associated with these strains and the challenges in managing them should not be underestimated. Further, the diversity of mobile genetic elements identified in both strains revealed their activity throughout evolutionary history. The whole genome sequencing method provided a better understanding of the genetic composition of *K. pneumoniae* KP-1 and KP-2. Furthermore, the differences in mobile genetic elements between the two strains highlight their varying activity during evolution.

The most striking suggestion is that the presence of numerous distinct CRISPR/Cas system regions and resistant genes in these isolates raises concerns about their potential impact on public health, the severity of the illnesses they can cause in humans, and the challenges in treating the associated consequences.

## Notes

### Competing Interest Statement

The authors have declared no competing interest.

